# MTSviewer: a database to visualize mitochondrial targeting sequences, cleavage sites, and mutations on protein structures

**DOI:** 10.1101/2021.11.25.470064

**Authors:** Andrew N. Bayne, Jing Dong, Saeid Amiri, Sali M.K. Farhan, Jean-François Trempe

**Affiliations:** Department of Pharmacology & Therapeutics and Centre de Recherche en Biologie Structurale, McGill University, Montréal, QC, Canada, H3G 1Y6; Department of Neurology and Neurosurgery, Montreal Neurological Institute, McGill University, Montréal, QC, Canada, H4A 3J1; Department of Human Genetics, McGill University, Montréal, QC, Canada, H4A 3J1

**Keywords:** MTSviewer, variant database, structure visualization, mitochondrial targeting signal, mitochondrial import, cleavage site, MTS

## Abstract

**Summary:** Mitochondrial dysfunction is implicated in a wide array of human diseases ranging from neurodegenerative disorders to cardiovascular defects. The coordinated localization and import of proteins into mitochondria are essential processes that ensure mitochondrial homeostasis and consequently cell survival. The localization and import of most mitochondrial proteins are driven by N-terminal mitochondrial targeting sequences (MTS’s), which interact with import machinery and are removed by the mitochondrial processing peptidase (MPP). The recent discovery of internal MTS’s - those which are distributed throughout a protein and act as import regulators or secondary MPP cleavage sites – has expanded the role of both MTS’s and MPP beyond conventional N-terminal regulatory pathways. Still, the global mutational landscape of MTS’s remains poorly characterized, both from genetic and structural perspectives. To this end, we have integrated a variety of tools into one harmonized R/Shiny database called MTSviewer (https://neurobioinfo.github.io/MTSvieweR/) which combines MTS predictions, cleavage sites, genetic variants, pathogenicity predictions, and N-terminomics data with structural visualization using AlphaFold models of human and yeast mitochondrial proteomes.

**Availability and Implementation:** MTSviewer is freely available on the web at https://neurobioinfo.github.io/MTSvieweR/.

Source code is available at https://github.com/neurobioinfo/MTSvieweR.

## 1. Introduction

Mitochondria are central to organismal health and regulate a diverse array of cellular processes, ranging from energy generation to immunity, proteostasis, and more (Mills et al. 2017; Ruan et al. 2017; Spinelli and Haigis 2018; Pfanner et al. 2019). Even though mitochondria contain their own genome, most mitochondrial proteins are nuclear encoded, translated in the cytosol, and imported into mitochondria (Wiedemann and Pfanner 2017). Consequently, mitochondria have evolved an intricate system of targeting and translocation to import these proteins through translocases of the outer (TOM) and inner (TIM) mitochondrial membranes, and sort them into their correct subcompartment (Neupert 2015). The most common targeting mechanism for matrix-localized proteins utilizes N-terminal mitochondrial targeting sequences (N-MTS), which form amphipathic helices and engage with TOM receptors before being passed through the TIM23 complex into the matrix (Callegari et al. 2020). In the matrix, N-MTS are cleaved off by the mitochondrial processing peptidase (MPP), which acts as a gatekeeper between import and overall mitochondrial quality control (Poveda-Huertes et al. 2017). The breadth of import mechanisms expands considerably when considering proteins localized to the intermembrane space (IMS), which typically lack an N-MTS and rely on disulfide trapping, or transmembrane (TM) proteins, which rely on a combination of accessory machinery and/or MTS’s for their insertion and sorting (Hansen and Herrmann 2019). It recently emerged that imported proteins can contain internal MTS’s (iMTS), which bind to TOM70 to regulate import rates and may also contain secondary MPP cleavage sites (Backes et al. 2018; Friedl et al. 2020). Furthermore, some proteins lacking an N-MTS still localize to and import into mitochondria via their iMTS’s (Bykov et al. 2022; Rahbani et al. 2021).

Mitochondrial targeting and import are innately linked to proteolysis, as mitochondria contain more than 40 proteases, coined “mitoproteases”, which regulate proteostasis, MTS removal, stress responses, signaling, and more (Deshwal et al. 2020). While MPP is the main protease implicated in N-MTS processing, other proteases act sequentially after MPP cleavage, including MIP, which removes an octapeptide, and XPNPEP3, which removes a single amino acid (Gomez‐Fabra Gala and Vögtle 2021). In specialized cases, other mitoproteases can regulate distal cleavages to drive signaling events, including PARL, a rhomboid protease which cleaves TM domains within the inner membrane (Spinazzi and de Strooper 2016; Lysyk et al. 2021). One example of a tandem MPP/PARL-cleaved protein is PINK1, a mitochondrial kinase that relies on its import and processing to either initiate or avoid the mitophagic cascade (Jin et al. 2010; Meissner et al. 2011; Bayne and Trempe 2019).

To facilitate the combined study of mitochondrial import and proteolysis, various tools have emerged, namely databases of mitochondrially localized proteins and prediction algorithms for sorting, MTS/iMTS propensity, and cleavage sites. In terms of mitoproteases, mass spectrometry experiments optimized for the labelling and enrichment of newly generated N-termini (neo-N-termini) have provided evidence for both canonical (ie. MTS removal) and non-canonical (ie. distal sites or N-terminal ragging) cleavage events within mitochondria (Calvo et al. 2017; Kleifeld et al. 2011; Vögtle et al. 2009). From a structural perspective, recent work has revealed the structures of human TOM and TIM complexes (Wang et al. 2020b; Qi et al. 2021), and of an iMTS-TOM70 complex between human TOM70 and the SARS-CoV2 protein ORF9b (Jiang et al. 2020). Still, how human MTS’s engage with and are passed across the other translocase subunits remain unclear. The structure of human MPP in complex with MTS substrates also remains unknown, which makes it difficult to confidently predict the consequences of MTS variants on MPP processing. From a genetic perspective, comparing the phenotypes of non-synonymous mutations within MTS’s, iMTS’s, or near cleavage sites may provide key insight into both areas, yet there is no database to facilitate this kind of analysis. There are also currently no resources to rapidly compare the outputs of the numerous mitochondrial prediction algorithms or to visualize MTS’s within 3D protein structures. To this end, we hope to expedite the genetic and structural interrogation of human mitochondrial proteins and their MTS’s with a novel database: MTSviewer (Fig. 1).

**Figure 1.**
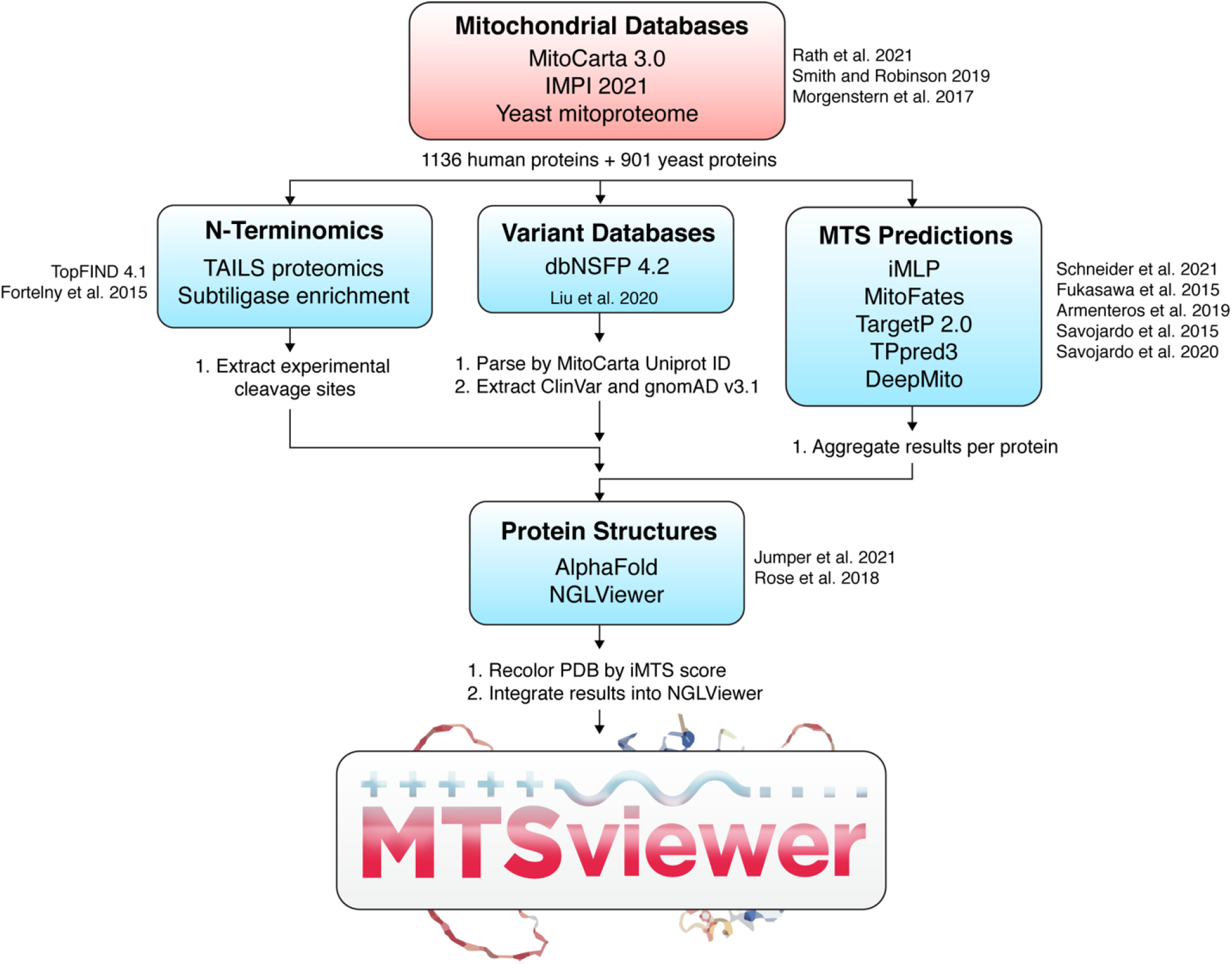
Workflow of MTSviewer. The database construction of MTSviewer, from initial mitochondrial databases to data integration and visualization.

## 2. Construction and content

The human mitochondrial proteome was downloaded from the MitoCarta 3.0 (1136 proteins) (Rath et al. 2021). Additional annotations for the MitoCarta protein list were appended from the Integrated Mitochondrial Protein Index (Q4pre-2021) (Smith and Robinson 2019). The yeast (*Sacchromyces cerevisiae*) mitochondrial proteome was derived from a high confidence dataset (901 proteins) (Morgenstern et al. 2017). Protein sequences were queried by UniProt ID and were submitted to: (1) iMLP – an internal MTS predictor using long short-term memory (LSTM) recurrent neural network architecture (Schneider et al. 2021); (2) TargetP2.0 – a presequence and cleavage site predictor using deep learning and bidirectional LSTM (Almagro Armenteros et al. 2019); (3) MitoFates – a presequence and cleavage site predictor using support vector machine (SVM) classifiers (Fukasawa et al. 2015); (4) TPpred3 – a targeting and cleavage site predictor using Grammatical Restrained Hidden Conditional Random Fields (Savojardo et al. 2015); (5) DeepMito – a sub-mitochondrial localization predictor using deep learning and convoluted neural networks (Savojardo et al. 2020). For cleavage sites derived from N-terminomics, mass spectrometry data were aggregated from TopFIND 4.1 by Uniprot ID of both human and yeast proteins (Fortelny et al. 2015). For variants and functional annotations of human proteins, dbNSFP v4.2a was parsed by Uniprot ID against GRCh38/hg38 coordinates (Liu et al. 2020). The resulting list was filtered using an in-house Python script into separate datasets for gnomAD v3.1 and ClinVar. Variants unique to the ExAC database were ignored. AlphaFold models for the *Homo sapiens* proteome (UP000005640) and *Sacchromyces cerevisiae* (UP000002311) were downloaded and matched by Uniprot ID (Jumper et al. 2021). An in-house Python script based on BioPandas (Raschka 2017) was used to parse the PDB files and re-color B-factors according to iMTS scores via iMLP. 3D visualization of protein structures was achieved using an adapted version of NGLViewer integrated into our R/Shiny application (Rose et al. 2018).

## 3. Utility and discussion

MTSviewer serves as a user-friendly platform for investigating MTS’s from both genetic and structural perspectives. The database requires minimal bioinformatics knowledge and features both human and yeast mitochondrial proteomes. With MTSviewer, users are able to: (1) compare mitochondrial prediction outputs from a variety of algorithms; (2) visualize MTS likelihood on a folded protein structure; (3) compare experimentally identified and predicted proteolytic events; (4) map non-synonymous variants (gnomAD, ClinVar, or user uploaded) within these MTS’s and cleavage sites. Using this platform, we have also curated a list of disease-linked variants within human MTS’s as a resource for their functional characterization.

## 4. User interface

The MTSviewer user interface is intuitive and begins by selecting or searching a gene of interest. Users specify the desired database for variant visualization (currently gnomAD v3.1 or ClinVar), and variants are overlaid onto an XY plot with the iMTS probability from protein N-to C-terminus. Hovering over a variant reveals cursory details which are fully expanded in the variant table. For the structure viewer, two coloring schemes are toggleable: the iMTS score, or the AlphaFold per-residue predicted local distance difference test (pLDDT) confidence score. Users can investigate specific residues or variants by clicking on the iMTS plot or 3D structure, and the structure viewer will automatically highlight the interactions (ie. polar contacts) and residues in proximity (5 Å) to the residue of interest (Fig. 2). Users can also upload custom variant lists for their proteins of interest in CSV format, which will be added to the iMTS propensity curve, populated into the variant list data tables, and become visualizable on the 3D protein structure. This feature allows users to compare where their variants lie in terms of MTS propensity, cleavage sites and other pathogenic variants on primary sequence and structural levels.

**Figure 2.**
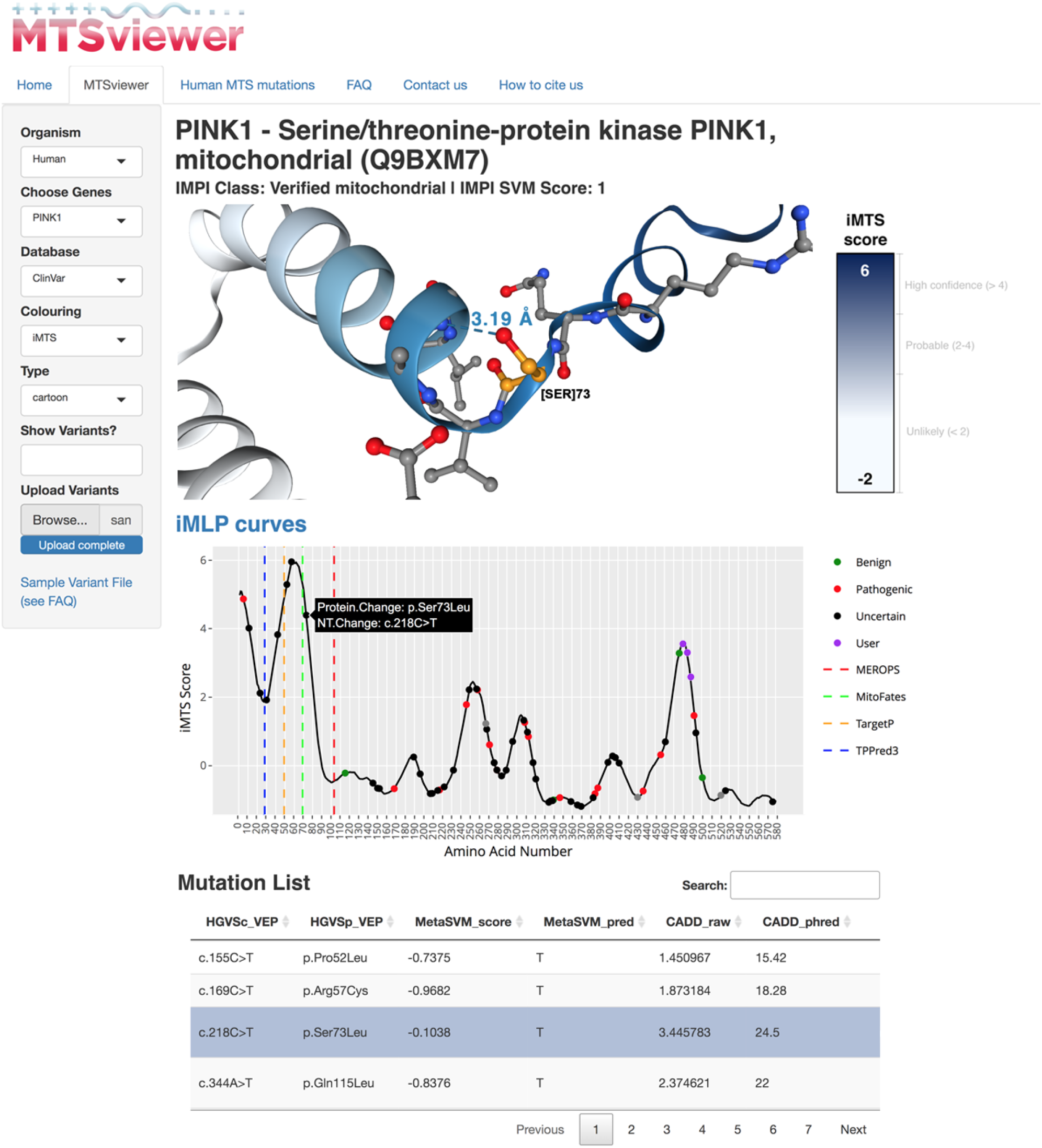
MTSviewer output for PINK1. A sample output from MTSviewer investigating human PINK1, a mitochondrial kinase with a uniquely long MTS and multiple predicted cleavage sites. Protein sequence, prediction algorithms, and N-terminomics data tables have been omitted for clarity but are available in full on the interactive MTSviewer web server. Ser73Leu has been highlighted as a variant of interest, as Ser73 is found within a region of high MTS propensity near the predicted MitoFates cleavage site.

The iMTS plot and structure viewer also contain toggleable visualizations to highlight cleavage site predictions from the various MTS predictors and/or experimentally determined N-terminomics sites. Aggregated comparisons of targeting predictors are pooled in table format, and data frames are exportable to facilitate downstream analyses. Taken together, these features enable users to rapidly generate protein-level hypotheses to test *in vitro*, or to rationalize previous *in vitro* findings with import- or protease-specific context.

## PINK1 as a case study

To highlight the utility of MTSviewer we have chosen PINK1 as a case study, given its cryptic N-MTS and the innate coupling of its import and processing to gate its accumulation on the TOM complex. Briefly, PINK1 is known to be cleaved by the rhomboid protease PARL in the IMM at Ala103, which is validated by the N-terminomics outputs seen in MTSviewer. The precise MPP cleavage site within the PINK1 N-MTS remains unknown, though an MPP-cleaved PINK1 fragment accumulates upon PARL knockdown (Greene et al. 2012). Based on the MTSviewer output for PINK1, there are many possibilities for the N-MTS MPP cleavage site, which will be critical to validate using *in vitro* assays, along with the effects of N-MTS variants (eg. Gly30Arg, Pro52Leu, Arg57Cys, and Ser73Leu). While some of these PINK1 N-MTS variants are still cleaved by PARL in healthy mitochondria and accumulate following mitochondrial damage (Sekine et al. 2019), their import rates and effects on MPP processing remain unstudied. Experiments which swap the PINK1 N-MTS with those from other mitochondrial proteins have shown that PINK1 can still be imported into mitochondria with chimeric N-MTS’s, though PINK1 accumulation is prevented (Kakade et al. 2022). While many of these N-MTS PINK1 chimeras can still be imported, their specific rates of import have also yet to be measured. This suggests that distal N-MTS elements of mitochondrial proteins (and variants within these regions) will be critical to study beyond the context of binary import success or blockage. Another useful feature of MTSviewer is the ability to gauge the length of a protein’s N-MTS by looking at the iMTS propensity plots. For reference, it has been estimated that MTS’s are usually 15-50 amino acids long (Wiedemann and Pfanner 2017), yet the PINK1 N-terminus exhibits high MTS propensity across its first 90 amino acids. As all of the MTSviewer iMTS data is available to download, users will be able to analyze global trends in MTS length and propensity across protein families to investigate the downstream consequences of longer or atypical N-MTS’s within mitochondrial proteins. Beyond the PINK1 N-MTS, the PINK1 iMTS plot within MTSviewer reveals a putative iMTS within the PINK1 C-terminus (a.a. 460-500), which could regulate PINK1 import or processing rates at the mitochondrial surface. It is known that PINK1 mRNA is co-transported with mitochondria (Harbauer et al. 2022), so it will be important to investigate the role of TOM70 binding to PINK1 and this putative iMTS during translation and import. The MTSviewer output for PINK1 also highlights the need to consider the oligomerization status of proteins when investigating their monomeric AlphaFold structures. PINK1 is known to dimerize on the OMM following depolarization which could occlude its iMTS in the folded dimeric state (Rasool et al. 2022; Okatsu et al. 2013), even if partially unfolded PINK1 monomers could bind to TOM70 upon import. Overall, MTSviewer will guide subsequent studies of atypically targeted proteins like PINK1 in the context of their MTS propensity, cleavage sites, and genetic variants.

## Comparison to similar databases

MTSviewer is the first interactive database to bridge genetic variants with mitochondrial targeting predictions, proteolytic evidence, and 3D protein structures. As such, it is essential to highlight the tools and databases that laid the foundation, and to highlight the gaps that our database aims to address. For MTS and MPP cleavage site predictions, TargetP2.0, MitoFates, and TPpred3 utilize orthogonal and sophisticated approaches, yet there remains no harmonized resource to compare their results. Our database currently only features these three predictors, as they are the most recently developed and performed best in benchmarking studies (Imai and Nakai 2020). For raw N-terminomics mass spectrometry data, TopFIND represents the gold standard for data accessibility and cleavage evidence across studies, but it does not provide genetic variants nor structural context for these proteolytic events (Fortelny et al. 2015). In terms of similar 3D structure viewers, the AlphaFold database contains its own module for visualizing contacts of a specified protein but does not allow for significant customizability (Jumper et al. 2021). ICN3D provides another alternative for user uploaded PDB visualization and manipulation, similar in complexity to the standalone PyMOL interface (Wang et al. 2020a). In terms of overall construction, MTSviewer resembles COSMIC-3D, which provides structural visualization for cancer genetics, with a specific focus on the druggability of protein targets (Jubb et al. 2018). KinaseMD has also taken a structural approach to the kinase mutational space, focusing on drug resistance, mutation hotspots, and network rewiring (Hu et al. 2021).

## Future developments and limitations

The current construction of MTSviewer features the inherent limitation that N-terminal MTS’s within AlphaFold predictions are typically low confidence and are depicted as unstructured. In the future, the inevitable structural determination of human MTS’s in complexes with TOM/TIM and/or MPP will enable us to model N-MTS’s more accurately and could be integrated as a scoring metric or docking module into later versions of MTSviewer. We will also implement a module for protease-specific exports (ie. variant lists near protease sites) to assess enrichment of pathogenic or uncharacterized variants near proteolytic sites. Overall, MTSviewer will be updated with new MTS prediction algorithms, experimental proteolytic evidence, and updated AlphaFold models on a regular basis.

## 4. Conclusions

MTSviewer is a novel R/Shiny database for investigating the mutational space, targeting sequences, proteolysis, and 3D structures of mitochondrial proteins. Users require minimal bioinformatics training and can rapidly generate variant lists, investigate structural consequences, compare the results of various mitochondrial prediction tools, and dissect potential cleavage sites.

## Declarations

### Ethics approval and consent to participate

Not applicable

### Consent for publication

Not applicable

### Availability of data and materials

The MTSviewer database is freely accessible via https://neurobioinfo.github.io/MTSvieweR/. Source code is available at https://github.com/neurobioinfo/MTSvieweR.

## Authors’ contributions

A.N.B. and J.F.T conceptualized MTSviewer. A.N.B., J.D., and S.A. created the original R/Shiny and Python codes used in database construction. All authors contributed to feature development, troubleshooting, and optimization of the database functionalities. A.N.B and J.F.T wrote the manuscript with contributions and editing from J.D., S.A., and S.F. All authors read and approved the final manuscript.

## Acknowledgements

We would like to thank Niels van der Velden for the ongoing support with NGLVieweR scripting and integration.

## References

Armenteros A, Emanuelsson SM, Winther O, Von Heijne O, Elofsson G. Detecting sequence signals in targeting peptides using deep learning. Life Science Alliance. 2019;2.

Backes S, Hess S, Boos F, Woellhaf MW, Gödel S, Jung M, et al. Tom70 enhances mitochondrial preprotein import efficiency by binding to internal targeting sequences. J Cell Biol. 2018;217:1369–82.

Bayne AN, Trempe J-F. Mechanisms of PINK1, ubiquitin and Parkin interactions in mitochondrial quality control and beyond. Cell Mol Life Sci. 2019;76:4589–611.

Bykov YS, Flohr T, Boos F, Zung N, Herrmann JM, Schuldiner M. Widespread use of unconventional targeting signals in mitochondrial ribosome proteins. EMBO J. 2022;41:e109519.

Callegari S, Cruz-Zaragoza LD, Rehling P. From TOM to the TIM23 complex -handing over of a precursor. Biol Chem. 2020;401:709–21.

Calvo SE, Julien O, Clauser KR, Shen H, Kamer KJ, Wells JA, et al. Comparative analysis of mitochondrial N-termini from mouse, human, and yeast. Mol Cell Proteomics. 2017;16:512–23.

Deshwal S, Fiedler KU, Langer T. Mitochondrial proteases: Multifaceted regulators of mitochondrial plasticity. Annu Rev Biochem. 2020;89:501–28.

Fortelny N, Yang S, Pavlidis P, Lange PF, Overall CM. Proteome TopFIND 3.0 with TopFINDer and PathFINDer: database and analysis tools for the association of protein termini to pre- and post-translational events. Nucleic Acids Res. 2015;43 Database issue:D290–7.

Friedl J, Knopp MR, Groh C, Paz E, Gould SB, Herrmann JM, et al. More than just a ticket canceller: the mitochondrial processing peptidase tailors complex precursor proteins at internal cleavage sites. Mol Biol Cell. 2020;31:2657–68.

Fukasawa Y, Tsuji J, Fu S-C, Tomii K, Horton P, Imai K. MitoFates: Improved Prediction of Mitochondrial Targeting Sequences and Their Cleavage Sites*. Molecular & Cellular Proteomics. 2015;14:1113–26.

Gomez-Fabra Gala M, Vögtle F-N. Mitochondrial proteases in human diseases. FEBS Lett. 2021;595:1205–22.

Greene AW, Grenier K, Aguileta MA, Muise S, Farazifard R, Haque ME, et al. Mitochondrial processing peptidase regulates PINK1 processing, import and Parkin recruitment. EMBO Rep. 2012;13:378–85.

Hansen KG, Herrmann JM. Transport of proteins into mitochondria. Protein J. 2019;38:330–42.

Harbauer AB, Hees JT, Wanderoy S, Segura I, Gibbs W, Cheng Y, et al. Neuronal mitochondria transport Pink1 mRNA via synaptojanin 2 to support local mitophagy. Neuron. 2022;110:1516-1531.e9.

Hu R, Xu H, Jia P, Zhao Z. KinaseMD: kinase mutations and drug response database. Nucleic Acids Res. 2021;49:D552–61.

Imai K, Nakai K. Tools for the recognition of sorting signals and the prediction of subcellular localization of proteins from their amino acid sequences. Front Genet. 2020;11:607812.

Jiang H-W, Zhang H-N, Meng Q-F, Xie J, Li Y, Chen H, et al. SARS-CoV-2 Orf9b suppresses type I interferon responses by targeting TOM70. Cell Mol Immunol. 2020;17:998–1000.

Jin SM, Lazarou M, Wang C, Kane LA, Narendra DP, Youle RJ. Mitochondrial membrane potential regulates PINK1 import and proteolytic destabilization by PARL. J Cell Biol. 2010;191:933–42.

Jubb HC, Saini HK, Verdonk ML, Forbes SA. COSMIC-3D provides structural perspectives on cancer genetics for drug discovery. Nat Genet. 2018;50:1200–2.

Jumper J, Evans R, Pritzel A, Green T, Figurnov M, Ronneberger O, et al. Highly accurate protein structure prediction with AlphaFold. Nature. 2021;596:583–9.

Kakade P, Ojha H, Raimi OG, Shaw A, Waddell AD, Ault JR, et al. Mapping of a Nterminal α-helix domain required for human PINK1 stabilization, Serine228 autophosphorylation and activation in cells. Open Biol. 2022;12:210264.

Kleifeld O, Doucet A, Prudova A, auf dem Keller U, Gioia M, Kizhakkedathu JN, et al. Identifying and quantifying proteolytic events and the natural N terminome by terminal amine isotopic labeling of substrates. Nat Protoc. 2011;6:1578–611.

Liu X, Li C, Mou C, Dong Y, Tu Y. dbNSFP v4: a comprehensive database of transcript-specific functional predictions and annotations for human nonsynonymous and splice-site SNVs. Genome Med. 2020;12:103.

Lysyk L, Brassard R, Arutyunova E, Siebert V, Jiang Z, Takyi E, et al. Insights into the catalytic properties of the mitochondrial rhomboid protease PARL. J Biol Chem. 2021;296:100383.

Meissner C, Lorenz H, Weihofen A, Selkoe DJ, Lemberg MK. The mitochondrial intramembrane protease PARL cleaves human Pink1 to regulate Pink1 trafficking: Rhomboid protease PARL cleaves human Pink1. J Neurochem. 2011;117:856–67.

Morgenstern M, Stiller SB, Lübbert P, Peikert CD, Dannenmaier S, Drepper F, et al. Definition of a high-confidence mitochondrial proteome at quantitative scale. Cell Rep. 2017;19:2836–52.

Mills EL, Kelly B, O’Neill LAJ. Mitochondria are the powerhouses of immunity. Nat Immunol. 2017;18:488–98.

Neupert W. A perspective on transport of proteins into mitochondria: a myriad of open questions. J Mol Biol. 2015;427 6 Pt A:1135–58.

Okatsu K, Uno M, Koyano F, Go E, Kimura M, Oka T, et al. A dimeric PINK1-containing complex on depolarized mitochondria stimulates Parkin recruitment. J Biol Chem. 2013;288:36372–84.

Pfanner N, Warscheid B, Wiedemann N. Mitochondrial proteins: from biogenesis to functional networks. Nat Rev Mol Cell Biol. 2019;20:267–84.

Poveda-Huertes D, Mulica P, Vögtle FN. The versatility of the mitochondrial presequence processing machinery: cleavage, quality control and turnover. Cell Tissue Res. 2017;367:73–81.

Qi L, Wang Q, Guan Z, Wu Y, Shen C, Hong S, et al. Cryo-EM structure of the human mitochondrial translocase TIM22 complex. Cell Res. 2021;31:369–72.

Rahbani JF, Roesler A, Hussain MF, Samborska B, Dykstra CB, Tsai L, et al. Creatine kinase B controls futile creatine cycling in thermogenic fat. Nature. 2021;590:480–5.

Raschka S. BioPandas: Working with molecular structures in pandas DataFrames. J Open Source Softw. 2017;2:279.

Rasool S, Veyron S, Soya N, Eldeeb MA, Lukacs GL, Fon EA, et al. Mechanism of PINK1 activation by autophosphorylation and insights into assembly on the TOM complex. Mol Cell. 2022;82:44-59.e6.

Rath S, Sharma R, Gupta R, Ast T, Chan C, Durham TJ. 0: an updated mitochondrial proteome now with sub-organelle localization and pathway annotations. Nucleic Acids Research. 2021;49:D1541–1547.

Rose AS, Bradley AR, Valasatava Y, Duarte JM, Prlic A, Rose PW. NGL viewer: webbased molecular graphics for large complexes. Bioinformatics. 2018;34:3755–8.

Ruan L, Zhou C, Jin E, Kucharavy A, Zhang Y, Wen Z, et al. Cytosolic proteostasis through importing of misfolded proteins into mitochondria. Nature. 2017;543:443–6.

Savojardo C, Bruciaferri N, Tartari G, Martelli PL, Casadio R. DeepMito: accurate prediction of protein sub-mitochondrial localization using convolutional neural networks. Bioinformatics. 2020;36:56–64.

Savojardo C, Martelli PL, Fariselli P, Casadio R. TPpred3 detects and discriminates mitochondrial and chloroplastic targeting peptides in eukaryotic proteins. Bioinformatics. 2015;31:3269–75.

Sekine S, Wang C, Sideris DP, Bunker E, Zhang Z, Youle RJ. Reciprocal roles of Tom7 and OMA1 during mitochondrial import and activation of PINK1. Mol Cell. 2019;73:1028-1043.e5.

Schneider K, Zimmer D, Nielsen H, Herrmann JM. Mühlhaus T. iMLP, a predictor for internal matrix targeting-like sequences in mitochondrial proteins. Biological Chemistry. 2021;402:937–43.

Smith AC, Robinson AJ. MitoMiner v4.0: an updated database of mitochondrial localization evidence, phenotypes and diseases. Nucleic Acids Res. 2019;47:D1225–8.

Spinazzi M, Strooper D. PARL: The mitochondrial rhomboid protease. Seminars in Cell & Developmental Biology. 2016;60:19–28.

Spinelli JB, Haigis MC. The multifaceted contributions of mitochondria to cellular metabolism. Nat Cell Biol. 2018;20:745–54.

Vögtle F-N, Wortelkamp S, Zahedi RP, Becker D, Leidhold C, Gevaert K, et al. Global analysis of the mitochondrial N-proteome identifies a processing peptidase critical for protein stability. Cell. 2009;139:428–39.

Wang J, Youkharibache P, Zhang D, Lanczycki CJ, Geer RC, Madej T, et al. iCn3D, a web-based 3D viewer for sharing 1D/2D/3D representations of biomolecular structures. Bioinformatics. 2020a;36:131–5.

Wang W, Chen X, Zhang L, Yi J, Ma Q, Yin J, et al. Atomic structure of human TOM core complex. Cell Discov. 2020b;6:67.

Wiedemann N, Pfanner N. Mitochondrial machineries for protein import and assembly. Annu Rev Biochem. 2017;86:685–714.

